# Rhythmic neuronal activities of the rat nucleus of the solitary tract are impaired by short-term high fat diet - implications for daily control of satiety by the brainstem

**DOI:** 10.1101/2021.03.02.433567

**Authors:** Lukasz Chrobok, Jasmin D Klich, Anna M Sanetra, Jagoda S Jeczmien-Lazur, Kamil Pradel, Katarzyna Palus-Chramiec, Mariusz Kepczynski, Hugh D Piggins, Marian H Lewandowski

## Abstract

Temporal partitioning of daily food intake is crucial for survival and involves the integration of internal circadian states and external influences such as the light-dark cycle and the composition of diet. These intrinsic and extrinsic factors are interdependent with misalignment of circadian rhythms promoting body weight gain, while consumption of a calorie dense diet elevates the risk of obesity and blunts circadian rhythms. Since cardiovascular disease, metabolic disorders, and cancer are comorbid with obesity, understanding the relationships between brain activity and diet is of pivotal importance. Recently, we defined for the first time the circadian properties of the dorsal vagal complex of the brainstem, a structure implicated in the control of food intake and autonomic tone, but if and how 24 h rhythms in this area are influenced by diet remains unresolved. Here we focused on a key structure of this complex, the nucleus of the solitary tract, and using a range of approaches, we interrogated how its neuronal and cellular rhythms are affected by high-fat diet. We report that short term consumption of this diet increases food intake during the day and blunts daily rhythms in gene expression and neuronal discharge in the nucleus of the solitary tract. These alterations in this structure occur without prominent body weight gain, suggesting that high-fat diet acts initially to reduce activity in the nucleus of the solitary tract, thereby disinhibiting mechanisms that suppress daytime feeding.

**GRAPHICAL ABSTRACT:** 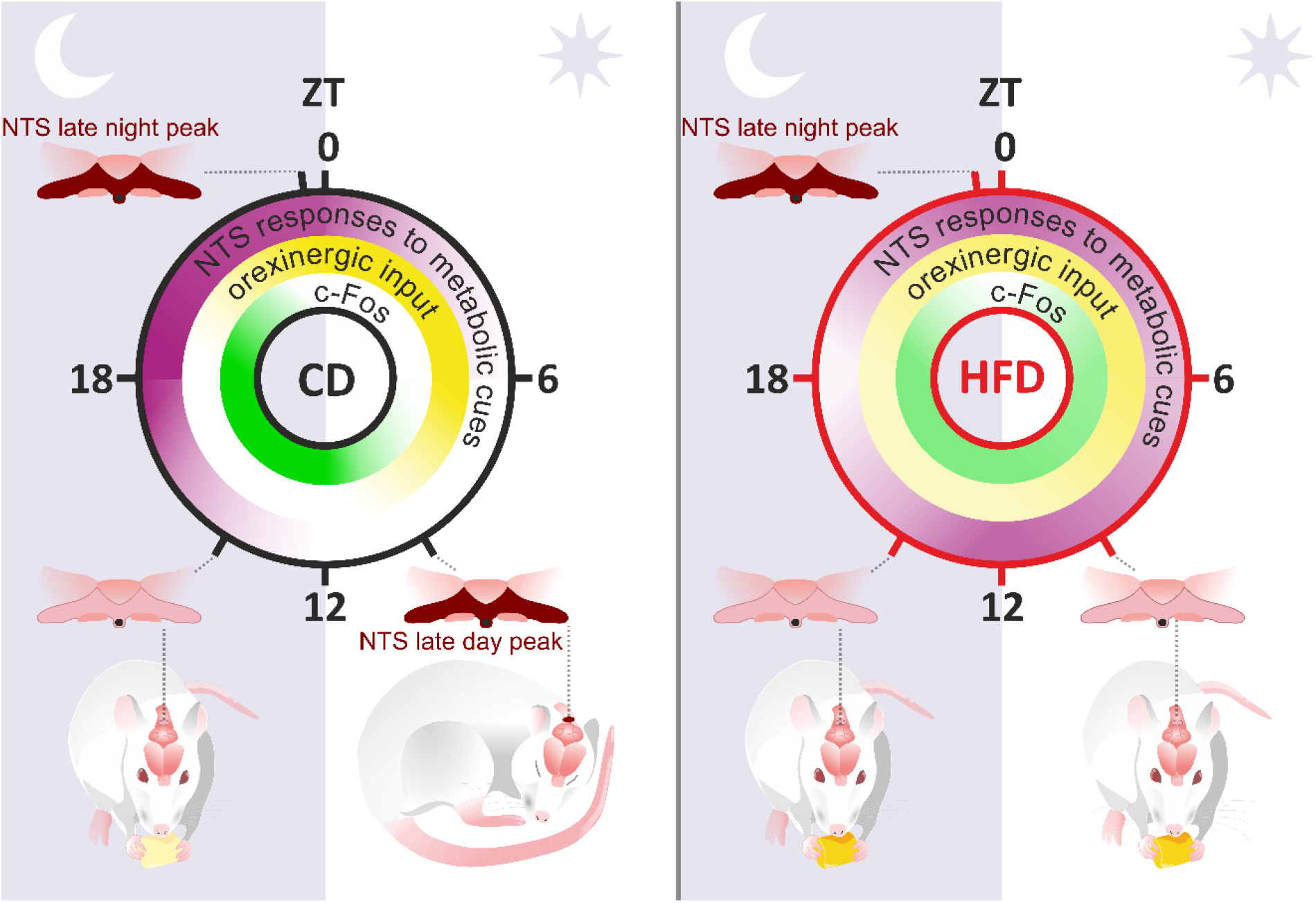

## 1. INTRODUCTION

Feeding is a vital and complex behaviour and precise control of meal initiation and termination is necessary for maintaining homeostasis and optimal health. Mammals have evolved specialised neural structures in the brainstem and hypothalamus to promote food seeking, its ingestion, and subsequent satiety and termination of feeding (Begg & Woods, 2013). Unfortunately, altering the constituents of a diet as well as timing of meals impedes the normal functioning of these neural mechanisms, leading to weight gain and comorbidities including metabolic syndrome, cardiovascular failure and some types of cancer (Vettor *et al*., 2002; Challet, 2015; Zarrinpar *et al*., 2016; Northeast *et al*., 2020*b*).

Most animals arrange their behavioural and physiological processes, including food seeking and ingestion, into preferred time of day or circadian phases – nocturnal rodents eat during the active dark phase and fast in behaviourally quiescent day (Gonnissen *et al*., 2013; Challet, 2015, 2019). Circadian timing is governed by circadian clocks, with the suprachiasmatic nuclei of the brain’s hypothalamus (SCN) functioning as the master timekeeper (Hastings *et al*., 2018). However, recent evidence questions the supremacy of the SCN, especially in the rhythmic control of feeding, with multiple extra-SCN oscillators expressing endogenous circadian properties independent of the master clock (Guilding & Piggins, 2007; Northeast *et al*., 2020*a*; Paul *et al*., 2020). To help conceptualise these findings, the term ‘food entrainable oscillator’ (FEO) has been coined in contrast with the SCN which is referred to as the ‘light entrainable oscillator’. Accumulating evidence demonstrates the FEO to be a network of peripheral tissues and metabolic cue processing brain sites, rather than a single nucleus orchestrating circadian rhythms in feeding behaviour (Verwey & Amir, 2009; Mistlberger & Antle, 2011; Mistlberger, 2011). Additionally, the appropriate phasing amongst tissue-specific oscillators is crucial for the expression of overall circadian rhythmicity, with their misalignment linked to metabolic and psychiatric disorders (Baron & Reid, 2014). Diet, body weight and optimal circadian rhythmicity are interlinked; long-term consumption of excessive calories and subsequent obesity negatively impact circadian processes, from the impairment of rhythmic clock genes expression to behavioural misalignment resulting in daytime feeding in nocturnal rodents (Kohsaka *et al*., 2007; Engin, 2017).

The dorsal vagal complex of the hindbrain (DVC) is a major hub for processing metabolic and cardiovascular cues and constitutes a brainstem satiety centre. The DVC can be divided into three tightly connected structures: (1) the area postrema (AP) – a sensory circumventricular organ detecting blood-borne information, (2) the nucleus of the solitary tract (NTS) – the major target of vagal afferents, processing both peripheral and central ingestive cues, and (3) the dorsal motor nucleus of the vagus (DMV) – an output part of the DVC populated by vagal motoneurons (Grill & Hayes, 2012). Recently, we reported robust intrinsic timekeeping properties of the mouse DVC with rhythmic clock gene expression (lasting up to several days in culture), neuronal activity and temporal regulation of blood-borne substances entry to the brain parenchyma (Chrobok *et al*., 2020). The DVC has also been linked to the development of obesity, with rhythms in core clock genes expression eliminated by long-term ingestion of a high-fat diet (HFD) (Kaneko *et al*., 2009). Indeed, the NTS is reported as a primary brain site at which long-term consumption of a HFD alters gene expression (Zhang *et al*., 2020).

Here, using a range of approaches, we interrogated at multiple levels how diet affects 24h patterns of food intake as well as daily variation in cellular and electrical activity of the rat NTS. We report that short-term consumption of a high-fat diet by nocturnal rats disrupts the daily pattern of ingestive activity by increasing daytime feeding. This is accompanied by blunted daily rhythms in NTS neuronal and cellular activity rhythms as well as the responsiveness of NTS neurons to metabolically relevant neuromodulators. These changes in behaviour and NTS molecular and cellular activity occur without prominent weight gain. From these findings, we speculate that a high-fat diet acts to dysregulate the NTS together with the daily patterning of food intake and that is these changes that precede excessive gain in body weight.

## 2. MATERIALS AND METHODS

### 2.1. Ethical approval

Experiments were approved by the Local (Krakow) Ethical Commission and performed in accordance with the European Community Council Directive of 24 November 1986 (86/0609/ EEC) and the Polish Animal Welfare Act of 23 May 2012 (82/2012). Every effort was made to reduce the number and suffering of animals used in the study.

### 2.2. Animals

All experiments were conducted on adult male Sprague Dawley rats. Animals were maintained in the Institute of Zoology and Biomedical Research animal facility at the Jagiellonian University in Krakow on a standard (12h/12h) light/dark cycle with water and food provided *ad libitum*. Constant environmental conditions (temperature: 23°C, humidity: ∼60%) were provided.

### 2.3. Diet

Rats were weaned at P28 and relocated from breeding colonies (with access to standard laboratory chow) and then randomly assigned to one of two dietary conditions. One group (CD) was fed a control diet (∼ 3,514 kcal/kg of chow, energy from: 10% fat, 24% protein, 66% carbohydrates; Altromin, Germany) and the other (HFD), a high-fat diet (∼ 5,389 kcal/kg of chow, energy from: 70% fat, 16% protein, 14% carbohydrates; Altromin). Rats used for electrophysiological experiments were maintained under these conditions for 2-3 weeks, whereas those used in immunohistochemical studies were culled following 4 weeks of either control or high-fat diet.

### 2.4. Food intake and body weight measurements

Male Sprague Dawley rats (n=19; 9 fed control diet and 10 fed high-fat diet) were individually housed in cages and then monitored throughout four weeks of diet. Body weight and food intake were measured at four time points across 24h (Zeitgeber Time: ZT0, 6, 12 and 18, where ZT12=lights-off and ZT0=lights-on) with initial measurements made three days following placement on the allocated diets (week 0) at P28. After four weeks of the diets, rats were culled at ZT6 (n=4 CD, n=5 HFD) and ZT18 (n=5/diet) by transcardial perfusion (see: *immunohistochemistry* section) and their stomachs collected and weighed.

### 2.5. Immunohistochemistry

#### 2.5.1. Tissue preparation and staining

A total of 50 male Sprague Dawley rats (24 fed control diet and 26 fed high-fat diet) were anaesthetised by isoflurane inhalation (2 ml/kg body weight; Baxter, USA) followed by an injection of sodium pentobarbital (100 mg/kg body weight, i.p.; Biowet, Poland). Next, tissue was fixed by transcardial perfusion with phosphate-buffered saline (PBS) followed by 4% paraformaldehyde (PFA) solution in PBS. Brains were removed and post-fixed in the same PFA solution overnight. Tissue preparation was performed at four time points across the 24 h light-dark cycle (starting at ZT0 with the interval of six hours). Six animals fed CD or HFD were culled per each time point (ZT0, 6, 12 and 18), except ZT 6 and 18 for HFD, when seven rats were killed.

Brain tissue was cut with a vibroslicer (Leica VT1000S, Germany) in 35 µm coronal slices. Three intersections along the anterior-posterior axis (rostral, intermediate and caudal) containing the DVC were collected. After sectioning, slices were moved to permeabilising solution containing 0.6% Triton-X100 (Sigma, Germany) and 10% normal donkey serum (NDS, Abcam, UK) in PBS at room temperature. After 35 min, slices were rinsed twice in PBS and incubated with primary antisera anti-OXB (raised in goat, 1:500; Santa Cruz Biotechnology, USA) and anti-NPY (raised in rabbit, 1:8000; Sigma) in PBS containing 2% NDS and 0.3% Triton-X100 overnight at 4°C. After that, slices were rinsed again and transferred to solution containing Cy3-conjugated secondary donkey anti-rabbit and AlexaFluor647-conjugated donkey anti-goat antisera (both 1:300, Jackson ImmunoResearch, USA) and incubated overnight at 4°C. Finally, sections were mounted on glass slides, dried and coverslipped with Fluoroshield^™^ (Sigma). Subsequently, slices were scanned using a Nikon A1-Si (Japan) confocal laser scanning system built on a Nikon Ti-E inverted microscope (Japan) at 20× magnification.

A separate sample of slices was stained with rabbit anti-cFos primary antisera (1:2000, Abcam) followed by secondary donkey anti-rabbit AlexaFluor 488 antibodies (1:300, Jackson ImmunoResearch) and scanned on a fluorescence microscope (Axio Imager.M2; Zeiss, Göttingen, Germany) under 20x magnification.

#### 2.5.2. Image analysis

Confocal images were further analysed in ImageJ software (NIH, USA). First, four Z-stack plains were combined into one image by maximal pixel intensity. Bernsen’s adaptive thresholding method was used to define regions of high local contrast. Next, circular regions of interest (ROIs) were outlined inside the anatomical boundaries of the NTS. NPY-ir in the DVC is the densest in the DMV, so it was used to assess NTS/DMV borders. The size of selection was adjusted so that six ROIs (three for each side) covered a greater part of the NTS. Area fraction (immunoreactive pixels/area of selection) was averaged from six measurements for every slice. The same settings were used to process all images. Nuclei stained against c-Fos were counted manually in the area of NTS in ZEN software (ZEN 2.3. black edition; Zeiss, Germany). For both OXB-ir and cFos, three sections (rostral, intermediate, and caudal) were analysed per rostrocaudal level of the NTS.

### 2.6. Electrophysiology

#### 2.6.1. Tissue preparation

Electrophysiological experiments were conducted on brain slices containing the dorsal vagal complex (DVC). Cull was performed at distinct time points across the 24h light-dark cycle (specific timing description in following sections). Animals were anaesthetised with isoflurane (2 ml/kg of body weight) and decapitated. Brains were quickly removed from the skull and placed in ice-cold preparation artificial cerebrospinal fluid (ACSF) composed of (in mM): 25 NaHCO_3_, 3 KCl, 1.2 Na_2_HPO_4_, 2 CaCl_2_, 10 MgCl_2_, 10 glucose, 125 sucrose with addition of pH indicator, Phenol Red 0.01 mg/l, osmolality ∼290 mOsmol/kg, continuously bubbled with carbogen (95% O_2_, 5% CO_2_). Tissue containing the brainstem was trimmed by the severing between the cerebellum and cerebral hemispheres and then mounted on the chilled plate of the vibroslicer (Leica VT1000S, Heidelberg, Germany) and cut into 250 μm-thick coronal slices. Those containing the DVC were placed in the pre-incubation chamber filled with carbogenated recording ACSF composed of (in mM): 125 NaCl, 25 NaHCO_3_, 3 KCl, 1.2 Na_2_HPO_4_, 2 CaCl_2_, 2 MgCl_2_, 5 glucose and 0.01 mg/l of Phenol Red (initial temperature: 32°C, cooled to room temperature) for at least one hour prior the recording.

#### 2.6.2. Multi-electrode array recordings

Multi-electrode array (MEA) experiments were performed at one of four time points across 24h on DVC slices obtained from 23 rats fed control or high fat diet (CD=11 and HFD=12 respectively) with a cull at (number of slices recorded for CD vs. HFD groups in brackets): ZT3 (n=4 vs. 4), ZT9 (n=5 vs. 5), ZT15 (n=4 vs. 5), ZT21 (n=4 vs. 5). After an incubation period, slices were positioned in the recording wells of the MEA2100-System (Multichannel Systems GmbH, Germany) with the NTS located just above a 6×10 recording array of perforated MEA (60pMEA100/30iR-Ti, Multichannel Systems) (Belle *et al*., 2021). Slices were perfused with carbogenated recording ACSF (2 ml/min) heated to 32°C throughout the experiment. After approximately half an hour of tissue settlement, recording was started (two hours after cull) and data acquired with Multi Channel Experimenter software (sampling frequency = 20 kHz; Multichannel Systems). After an hour of baseline recording, drugs were diluted in 6 ml of the recording ACSF to a final concentration of 200 nM (OXA) or 1 µM (GLP-1) and were administered by bath perfusion in one hour intervals.

Additionally, spontaneous firing activity was evaluated by 36h-long recordings of eleven DVC slices obtained from three rats fed control diet and three fed high-fat diet. To preserve firing activity and good condition of recorded tissue, these recordings were performed in the ACSF enriched with 1 mg/ml penicillin-streptomycin (Sigma) at 25°C with additionally reduced suction. Data was collected for 1 min in 10 min intervals and merged after recording with custom made MatLab script (R2018a version, MathWorks, USA). For this set of experiments, animals were culled at ZT0 and recordings started at ZT2, with five slices obtained from high-fat fed and six from control diet fed rats.

#### 2.6.3. Drugs

Orexin A (OXA; 200 nM, Bachem, Switzerland) and glucagon-like peptide 1 (GLP-1; 1µM, Bachem) were stocked at 100× concentration at -20°C and were freshly diluted in the recording ACSF prior the application by bath perfusion.

#### 2.6.4. Spike-sorting and analysis of multi-electrode array data

Raw data was exported to HDF5 files with Multi Channel DataManager (Multichannel Systems GmbH) and processed through a custom-made MatLab script (R2018a version, MathWorks) to remap and convert the file to DAT format. DAT files were initially automatically spike-sorted with KiloSort program (Pachitariu *et al*., 2016) in MatLab environment. To enhance the speed of spike-sorting, a GPU was used (NVIDIA GeForce GTX 1050Ti GPU; CUDA 9.0 for Windows). In parallel, raw data was exported to CED-64 files with MultiChannel DataManager, and subsequently remapped and filtered with Butterworth band pass filter (fourth order) from 0.3 to 7.5 kHz by a custom-made Spike2 script. Spike-sorting results were transferred into the prepared CED-64 files (Spike2 8.11; Cambridge Electronic Design Ltd.) for further visualisation using a custom-written MatLab script. Each putative single-unit automatically sorted by KiloSort was then verified in Spike2 8.11 with the aid of autocorrelation, principal component analysis (PCA) and spike shape inspection. If the autocorrelation contained unfeasibly brief inter-spike intervals (shorter than a refractory period) and/or the PCA revealed distinct clustering, these spike-sorting results were further manually refined.

Baseline comparisons for short-term recordings were made from 1800 s of spontaneous firing at the beginning of the protocol. Reactions to drug applications were measured in 30 s bins with NeuroExplorer 6 (Nex Technologies, USA). If the single-unit activity (SUA) varied by three standard deviations (SDs) from the baseline values after the drug application, the unit was deemed responsive. Response amplitudes were further calculated as a difference between 600 s baseline mean frequency and maximal value during the response to a drug.

Long term recordings were analysed in 1 min bins as electrophysiological activity was sampled for 1 min every 10 min. In order to avoid data misinterpretation caused by tissue degradation, only the first 30 h of each recording was analysed. Time of peak firing was manually selected. The unit was deemed ‘rhythmic’ if its spontaneous activity throughout the 30h revealed a single predominant peak. Those units whose firing rate did not change throughout the recording or displayed multiple peaks were classified as ‘non-rhythmic’. Activity heatmaps were prepared from multi-unit and single-unit activity (MUA and SUA, respectively) with custom made MatLab script (R2018a version, MathWorks, USA). Additionally, heatmaps presenting SUA over time were depicted on smoothed data (Gaussian filter, width: 5 bins) as the firing normalised for each unit separately (maximal firing rate = 1), with units sorted by the time of manually depicted peaks. Non-rhythmic units were placed below in a random order. Time of peak histograms were calculated in 1 h bins. Rayleigh plots of peak phase and subsequent r values were calculated in El Temps (University of Barcelona, Spain). For these plots, circles represent peak firing rate for single units, the arrow indicates the phase vector and its length represents significance with respect to the inner dotted circle of the *p*=0.05 threshold. Variance is represented by the black box around arrow heads. Average time of peak activity was calculated as circular means with the use of Circular Statistics toolbox in MatLab (R2018a version, MathWorks).

### 2.7. Statistics

Statistical testing was generally performed in Prism 7 (GraphPad). *P* values <0.05 were considered significant. All data in the manuscript text is presented as mean ± SD.

Repeated measures (RM) two-way ANOVA was used to assess time and diet-dependent changes in (1) body weight, (2) calorie intake, and (3) calorie intake at day or night phases. Further RM ANOVA was used to compare circadian waveforms of the spontaneous neuronal activity in long-term MEA recordings. When RM two-way ANOVA was used, individual means were assessed by Dunn’s multiple comparison test.

Independent two-way ANOVA was performed to evaluate significance of daily and diet-dependent differences in (1) stomach weight, (2) number of cFos+ cells, (3) density of OXB-ir fibres, (4) spontaneous neuronal activity in acute MEA recordings, and (5) amplitude of responses to metabolically relevant peptides in these recordings. Subsequent post-hoc multiple comparison of individual means was performed with Sidak’s test.

Modality of distributions was assessed by Hartigan’s dip test in MatLab (R2018a version, MathWorks), where *p*<0.05 pinpointed significantly multimodal distribution. Circular means were compared using the circular analogue of ANOVA (Watson-Williams test) performed with Circular Statistics toolbox in MatLab (R2018a version, MathWorks).

## 3. RESULTS

### 3.1. Short-term high-fat diet alters daily feeding pattern

Previous reports clearly demonstrate a reciprocal relation between obesity and circadian rhythmicity: circadian misalignment promotes obesity, and conversely, calorie-dense diets disturb a plethora of circadian processes (Zarrinpar *et al*., 2016; Engin, 2017). Here we used rats to address whether short-term consumption of a high-fat diet changes their daily pattern of ingestion and if so, whether this was accompanied by excessive weight gain. Therefore, we fed rats either control or a high-fat diet and profiled feeding behaviour and body weight at four time points of the day over a four week test period. The first measurement (week 0) was made three days after the start of the protocol, when rats (P28) had been randomly assigned to two groups: an experimental group (HFD) fed a high-fat diet (70% kcal from fat, n=10), and a control group (CD) fed a matched control diet (10% kcal derived from fat, n=9). Throughout the experiment, rats exhibited a significant time of day variation in body weight (ZT: *p*<0.0001, RM two-way ANOVA; Fig. 1A), however after three weeks, this daily pattern became blunted in the HFD rats (week 3: *p*=0.0110, week 4: *p*=0.0023, RM two-way ANOVA interaction; Fig. 1A). This was evident after four weeks of diet, when CD rats displayed an early day weight loss (seen throughout all weeks of our protocol), and this was significantly reduced in the HFD group (amplitude from ZT0 to ZT6 at week 4: *p*=0.0003, unpaired t-test; Fig. 1A). This change in the daily pattern of feeding was not attributable to group differences in body weight gain since throughout the course of the experiment, high-fat fed rats did not significantly differ in body weight from CD animals (diet: *p*=0.5563, RM two-way ANOVA; Fig. 1C).

**Figure 1.**
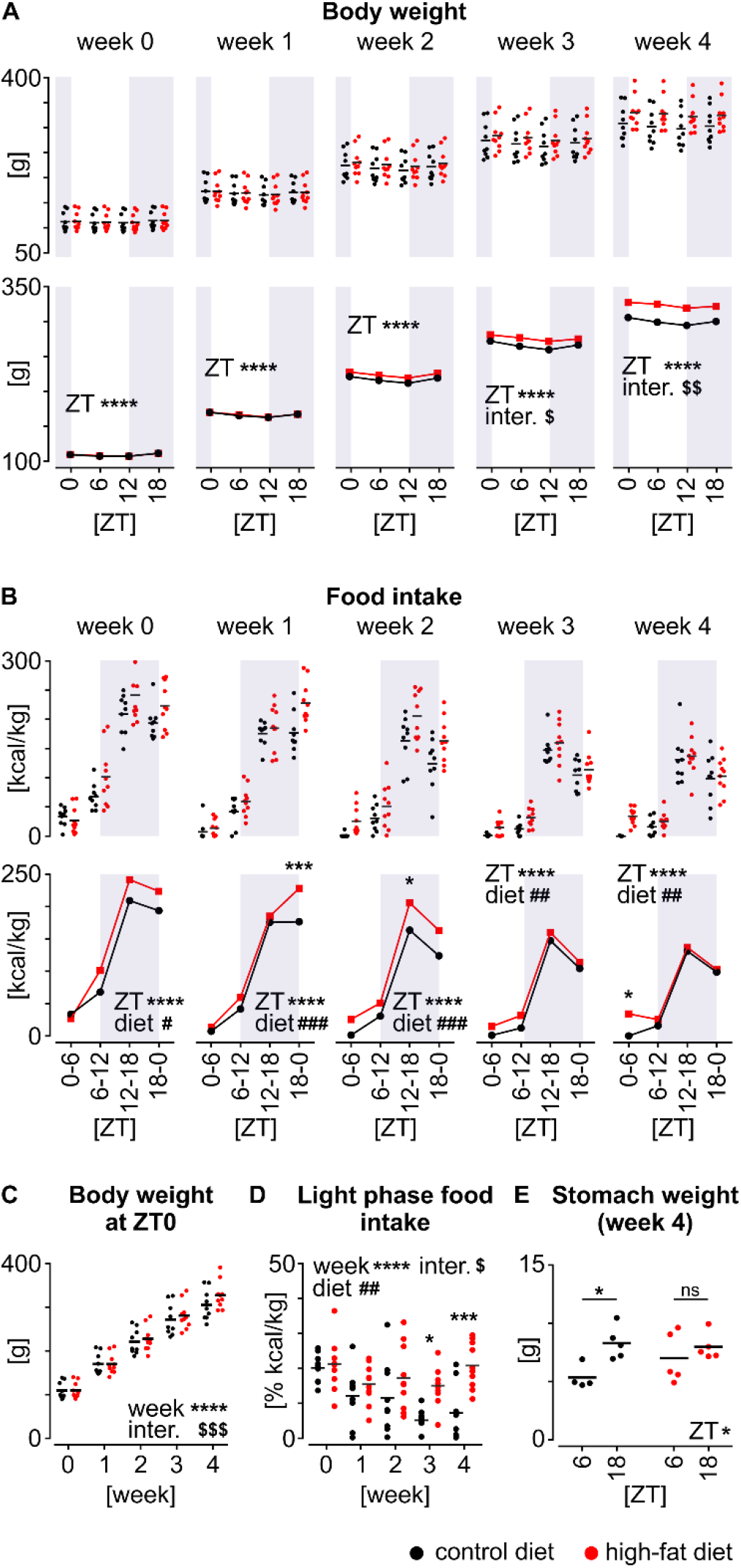
Short term high fat diet (HFD) reorganises the daily partitioning of feeding with increased daytime feeding preceding excessive body weight gain. (**A**) Daily variation in body weight over the four weeks of diet (HFD: n=10; control diet, CD: n=9). Note the progressively changing shapes of these profiles for rats fed HFD (ZT: \*\*\*\**p*<0.0001, RM two-way ANOVAs; $*p*=0.0110, $$*p*=0.0023, RM two-way ANOVA interactions). Top panel shows individual measurements with means, whereas mean profiles are drawn in the bottom panel. (**B**) Matched daily measurements of food intake in 6 h bins, calculated as kcal per kg body weight (ZT: \*\*\*\**p*<0.0001, diet0: #*p*=0.0142, diet1: ###*p*=0.0008, diet2: ###*p*=0.0002, diet3: ##*p*=0.0063, diet4: ##*p*=0.0095, RM two-way ANOVAs). Asterisks adjacent to plots mark results of multiple comparison test (week1: \*\*\**p*=0.0007, week2: \**p*=0.0393, week4 \**p*=0.0369, Sidak’s tests). Note the switch to diurnal overeating in animals given HFD. (for *A* and *B*) Top panels show individual measurements with means, whereas mean profiles are drawn in the bottom panels. (**C**) Increase in body weight over four weeks of the experiment, faster for HFD-fed rats (week: \*\*\*\**p*<0.0001, interaction: $$$*p*=0.0003, RM two-way ANOVA). (**D**) Proportion of the food intake during the resting light phase, calculated as % of total kcal per kg body weight (week: \*\*\*\**p*<0.0001, diet: ##*p*=0.0037, interaction: $*p*=0.0220, RM two-way ANOVA). Animals fed HFD significantly increase diurnal feeding in the third (\**p*=0.0153) and fourth week of diet (\*\*\**p*=0.0004, Sidak’s test). (**E**) Stomach weight of rats culled after four weeks of diet in ZT6 (n=9) and ZT18 (n=10). Note the day-to-night difference in the CD only (ZT: \**p*=0.0143, two-way ANOVA; CD: \**p*=0.0235, HFD: ns *p*=0.5541, Sidak’s test). In all graphs, red represents HFD, whereas black CD.

Throughout the four weeks of assessment, HFD rats consumed more calories per 24 h than CD animals (diet: week 0: *p*=0.0142, week 1: *p*=0.0008, week 2: *p*=0.0002, week 3: *p*=0.0063, week 4: *p*=0.0095, RM two-way ANOVAs; Fig. 1B). The day-night partitioning of food intake also varied in a diet-related manner. Initially, HFD rats increased their nocturnal food intake, but after three weeks on this diet, these animals gradually shifted to consume excessive calories during the light phase (food intake from ZT0 to ZT6 at week 4: *p*=0.0369, Sidak’s test; Fig. 1B). Indeed, following four weeks on the different diets, high-fat fed animals consumed 20.6 ± 2 % of their daily calories during the lights-on phase, whereas CD rats consumed only 7.1 ± 3 % of daily calories during this phase (*p*=0.0004, Sidak’s test; Fig. 1D). Consistent with this altered pattern of day-night feeding, assessment of stomach weight of animals culled at the mid-day (ZT6) or mid-night (ZT18) point on the last day of week 4 of the experiment showed a time of day effect (ZT: *p*=0.0143, two-way ANOVA; Fig. 1E). In CD rats, stomach weight increased significantly from ZT6 to ZT18 (*p*=0.0235), whereas, there was no difference in HFD animals (*p*=0.5541, Sidak’s tests; Fig. 1E; statistical details summarised in Supplementary Table 1). These findings indicate that prior to excess gain in body weight, rats respond to the high fat diet by altering their daily partitioning of calorie intake through increasing food consumption during the day.

### 3.2. High-fat diet shifts and blunts daily cFos expression in the NTS

Evidence from a recent study localises gene expression changes evoked by long-term HFD predominately to two metabolic information processing areas in the central nervous system: the dorsomedial hypothalamus and the NTS (Zhang *et al*., 2020). Stomach distension elevates the expression of cFos, a proxy of neuronal input/activity in the NTS (Sabbatini *et al*., 2004; Roelofs *et al*., 2020) and to investigate how short-term consumption of a high-fat diet influences daily patterns of expression of cFos in the NTS, we gave a new cohort of rats the high-fat diet (n=26; HFD group) or the control diet (n=24; CD group). After four weeks of *ad libitum* consumption of these diets, animals were culled at one of four time points (ZT0, 6, 12 and 18; n=6-7 rats/time point/diet) and cFos expression was examined at three distinct rostrocaudal levels of the coronally sectioned NTS: rostral (immediately rostral to the area postrema, AP), intermediate (with AP) and caudal (at the level of the obex). Time of day variation in the number of cFos-positive cells in the NTS was evident in all sections, with low day-time and high night-time values (ZT: *p*<0.0001, two-way ANOVA; Fig. 2A-C).

**Figure 2.**
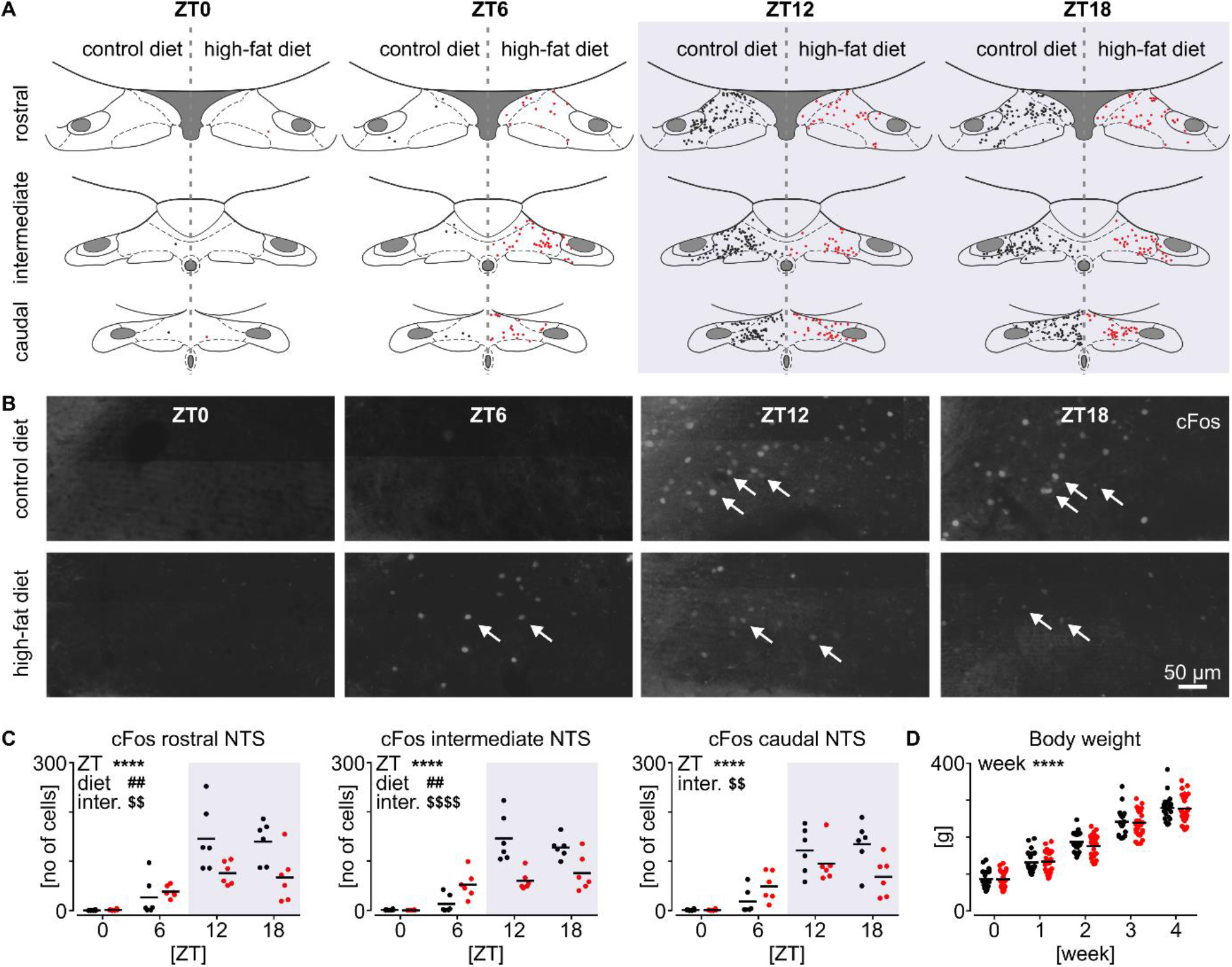
Four weeks of high fat diet (HFD) disrupts daily variation in cFos expression. (**A**) Schematic representation of cFos-positive cells in three sections of the nucleus of the solitary tract (NTS, black and red circles) at four time points across 24h (ZT0,6,12,18). At each section, left side represents control diet (CD) and right, results obtained from high-fat diet (HFD) animals. (**B**) Example epifluorescence images from the centre of the NTS. White arrows point individual cFos+ nuclei. (**C**) Daily profiles showing change in the number of cFos-positive cells in three sections of the NTS: rostral (ZT: \*\*\*\**p*<0.0001, diet: ##*p*=0.0041, interaction: $$*p*=0.0083), intermediate (ZT: \*\*\*\**p*<0.0001, diet: ##*p*=0.0023, interaction: $$$$*p*<0.0001) and caudal (ZT: \*\*\*\**p*<0.0001, interaction: $$*p*=0.0088, two-way ANOVAs). (**D**) Changes in body weight over four weeks of diet in this cohort of animals (week: \*\*\*\**p*<0.0001, RM two-way ANOVA). In all plots, red represents HFD, whereas black CD, and n=6-7 rats/ZT/diet.

This variation was significantly affected by HFD, which decreased overall numbers of cFos-positive cells in intermediate (diet: *p*=0.0023) and rostral NTS sections (diet: *p*=0.0041, two-way ANOVAs; Fig. 2A-C); the levels of the NTS implicated in processing metabolic information (Grill & Hayes, 2012). Intriguingly, the shape of the 24h profile of cFos expression in the NTS was also altered by high-fat diet, being damped during the night and elevated around the middle of the day in HFD animals. This was apparent in the rostral (*p*=0.0083) and caudal (*p*=0.0088) and was most prominent in the intermediate NTS (*p*<0.0001, two-way ANOVA interactions; Fig. 2A-C). Consequently, the middle day to early night increase in cFos observed at all NTS levels in CD rats (rostral: *p*<0.0001, intermediate: *p*<0.0001, caudal: *p*<0.0001; Sidak’s tests) was absent in HFD animals (rostral: *p*=0.4229, intermediate: *p*=0.9966, caudal: *p*=0.1269; Sidak’s tests). It would be tempting to attribute the absence of such daily variation in NTS cFos of HFD rats to excessive weight gain, but no significant change in body mass was found (diet: *p*=0.7182, RM two-way ANOVA; Fig. 2D). Since the increase in NTS cFos of rats fed the control diet occurs as the animals transition from the behaviourally quiescent day to the active night-time, this likely reflects the intrinsic daily rhythm of NTS neurons and/or feedback from stomach distension resulting from the initial feeding bouts. In summary, our findings show that the daily pattern of cFos expression in the NTS is significantly modified by high-fat diet; in comparison to control diet fed rats, cFos expression is increased during the day and decreased at night. These changes in NTS cellular activity occur without prominent change in body weight, but do parallel the altered daily feeding pattern of HFD rats noted in outlined above section 3.1. Summary of statistical analysis of this dataset can be found in Supplementary Table 2.

### 3.3. Orexinergic input to the NTS is compromised by high-fat diet

In rodents, the expression of hypothalamic neuropeptides implicated in metabolic control and energy homeostasis is downregulated through long-term consumption of a HFD (Hariri & Thibault, 2010; Velloso & Schwartz, 2011). To investigate whether brainstem input from the appetite-promoting orexin neurons of the lateral hypothalamus is altered by diet, we assessed orexin B (OXB)-immunoreactivity (-ir) in brainstem sections obtained from the same cohort of 26 HFD and 24 CD-fed rats used above to examine cFos expression at the 6h intervals over the day-night cycle. We found that the density of OXB-ir fibres exhibited time of day variation (ZT: *p*=0.0259), with higher levels during the daytime, but only in the intermediate level of the NTS. No time-dependent variation was present at the caudal (ZT: *p*=0.3341) or rostral levels (ZT: *p*=0.0936; two-way ANOVAs; Fig. 3A,B). Consistent with alteration in hypothalamic neuropeptide expression observed in HFD mice (Nobunaga *et al*., 2014), here in the rat, high-fat diet significantly decreased OXB-ir at the intermediate (diet: *p*=0.0051) and caudal levels (diet: *p*=0.0215) of the NTS, and approached significance in the rostral NTS (diet: *p*=0.0936, two-way ANOVAs; Fig. 3A,B). This suggests that daily variation in orexin input to the NTS is suppressed as a consequence of consumption of the calorie dense diet, with the intermediate level of the NTS being most affected by diet. This raises the possibility that excess fat consumption can act at an origin of NTS afferents to regulate an important neural input. Statistical recapitulation is depicted by Supplementary Table 3.

**Figure 3.**
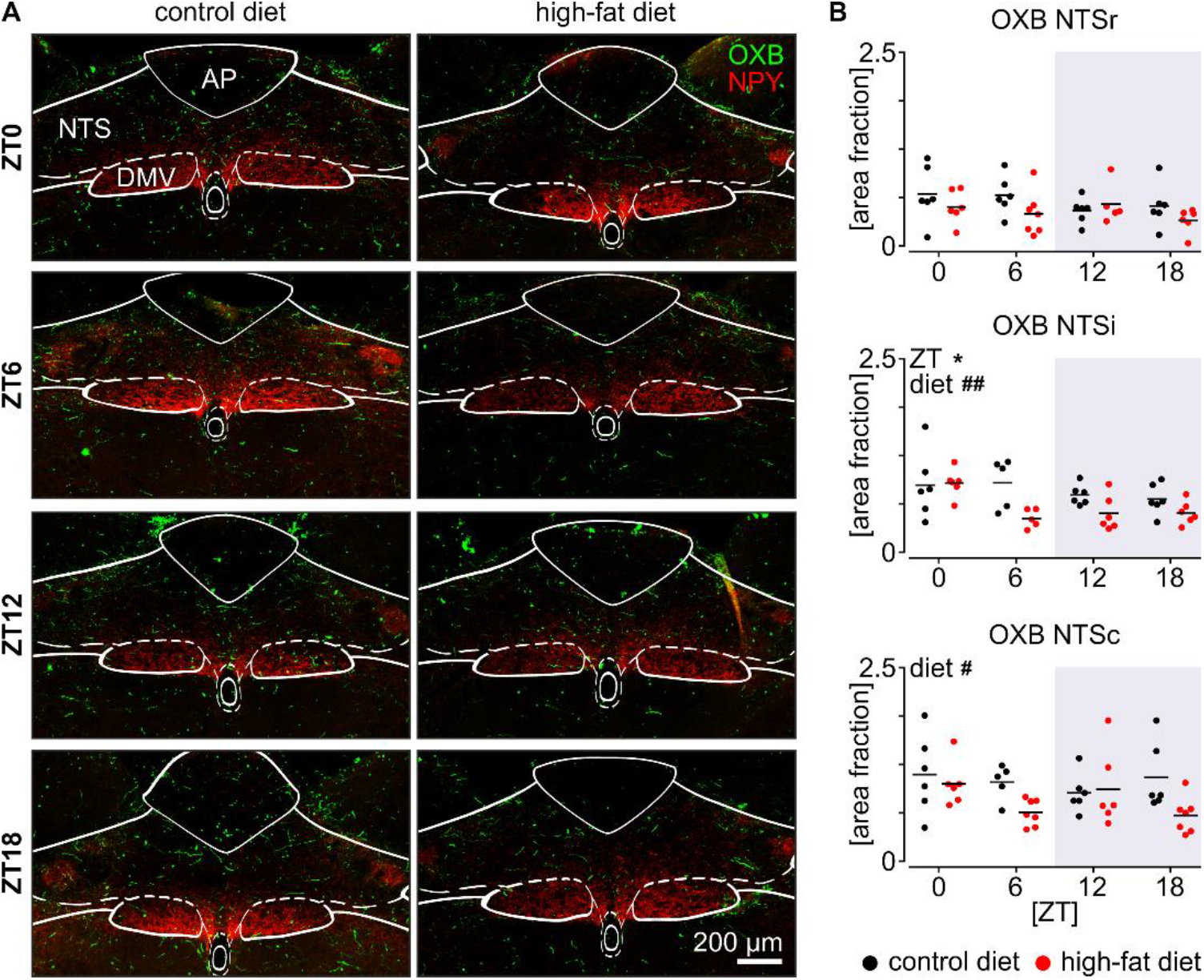
Short-term high fat diet (HFD) downregulates orexinergic immunoreactivity in the nucleus of the solitary tract (NTS). (**A**) Example confocal photomicrographs depicting orexin B (OXB)-ir fibres and neuropeptide Y(NPY)-ir fibres in green and red, respectively, with borders of the dorsal vagal complex components outlined: AP – area postrema, DMV – dorsal motor nucleus of the vagus. (**B**) Daily changes of OXB-ir fibre density displayed as area fraction for the three intersections of the NTS: rostral (NTSr), intermediate (NTSi; ZT: \**p*=0.0259, diet: ##*p*=0.0051) and caudal (NTSc; diet: #*p*=0.0215). In all plots, red represents HFD, whereas black CD, and n=6-7 rats/ZT/diet.

### 3.4. Short-term high-fat diet delays endogenous circadian rhythm of the DVC *ex vivo*

The elevated daytime feeding pattern of rats fed HFD may arise from shifted activity of the DVC or the recurrent daytime distension of the stomach caused by feeding may feedback to re-model DVC neuronal activity (Sabbatini *et al*., 2004). To gain insight into the direction of this causality, we performed electrophysiological recordings of the DVC with a multi-electrode array (MEA) platform in two protocols: (1) to assess changes in NTS physiology arising from intrinsic time of day processes as well as feedback from behavioural challenges, acute short-term recordings of neural activity were made 2h after cull at four time points across 24h; and (2) to evaluate intrinsic circadian regulation of DVC physiology alone, long-term (up to 36h) recordings of spontaneous firing of DVC neurons were initiated 2h after cull at ZT0.

For the first protocol, we initially recorded 30 min of spontaneous multi-unit neuronal activity from 36 acute DVC brain slices (17 from rats fed CD and 19 – HFD) and further spike-sorted single units obtained from electrodes located in the NTS (Fig. 4A,B). Animals were culled at ZT3, 9, 15, and 21, with recordings always begun 2h later (ZT5, 11, 17 and 23). Time of day variation in neuronal activity was evident in the NTS (ZT: *p*=0.0056, two-way ANOVA), which was significantly altered by HFD (*p*<0.0001; two-way ANOVA interaction) through increasing NTS neuronal activity (diet: *p*=0.0001, two-way ANOVA; Fig. 4C). In rats fed the control diet, NTS neurons greatly increase their activity from ZT5 to a zenith near the end of light phase (ZT11) (CD, *p*<0.0001). This elevation is eliminated in rats fed the high-fat diet (HFD, *p*=0.9998, Sidak’s tests), with the higher level of activity occurring during the late night (ZT23). These findings indicate that NTS neurons increase their activity from early to late day and that consumption of a HFD blunts this and delays peak activity to later in the night. The possibility remains however that such diet-related changes may be a consequence of abnormal feeding behaviour. Results of statistical analysis of this section can be found at Supplementary Table 4.

**Figure 4.**
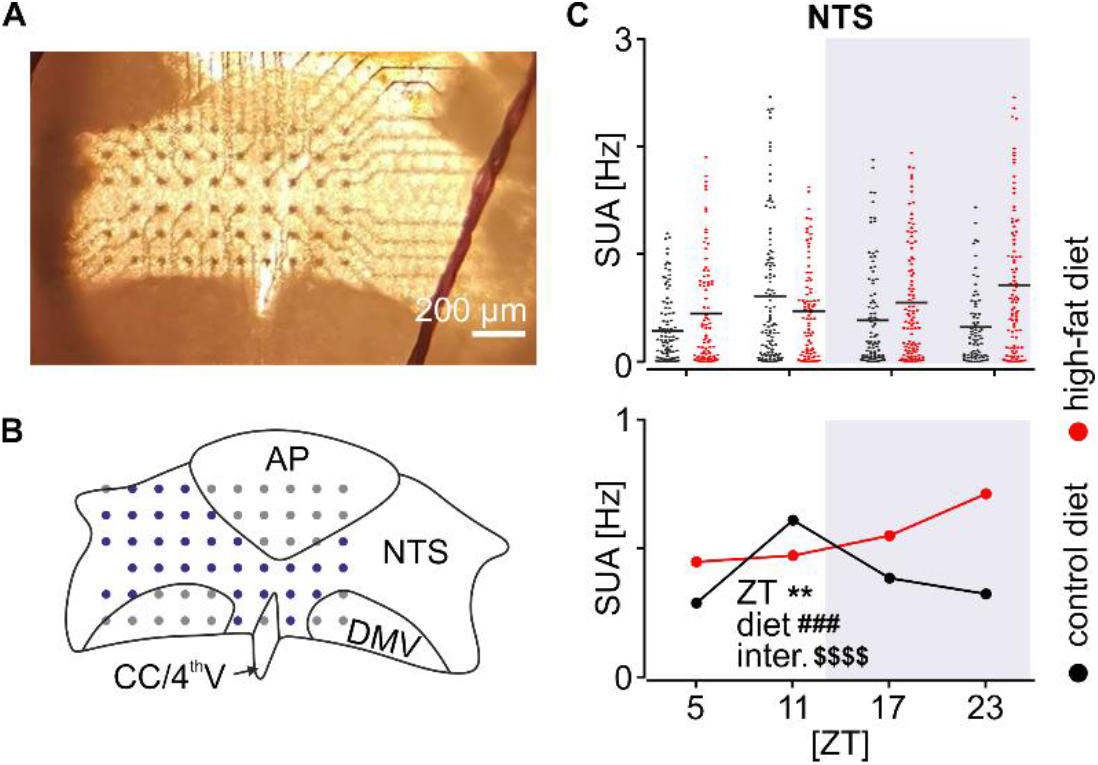
Daily patterning of neuronal activity in the nucleus of the solitary tract (NTS) is modified by short-term high-fat diet (HFD). (**A**) Representative image of the DVC slice upon the multi-electrode array (MEA). (**B**) Reconstruction of the recording locations with the outline of DVC areas: AP – area postrema, CC/4^th^V – central canal/4^th^ ventricle, DMV – dorsal motor nucleus of the vagus, NTS – nucleus of the solitary tract. Electrodes located in the NTS are coloured in blue. (**C**) Scatterplot of individual values (*above*) and a summary plot (*below*) of the single-unit activity in the NTS at four time points across 24h: ZT5, 11, 17 and 23 (ZT \*\**p*=0.0056, diet ###*p*=0.0001, $$$$interaction *p*<0.0001; two way-ANOVA). Data were shown as mean. In plots, red codes high-fat diet, and black – control diet.

To directly assess how diet influences the intrinsic circadian regulation of DVC neuronal activity, we made 36h-long recordings of the spontaneous neuronal activity throughout the DVC in brainstem slices prepared at ZT0 from three rats fed CD (six slices) and three rats fed HFD (five slices). These investigations enabled us to assess not only neuronal discharge in the NTS, but also spontaneous electrophysiological activity in the DMV as well as small number of neurons in the AP. To gain additional insight into the circadian phasing of the DVC neuronal activities, in these recordings we also studied the spontaneous activity of neurons in the DMV, and a small number of neurons in the AP. All recordings were initiated at pZT2 (projected ZT) and the first 30 h of these recordings was subsequently analysed. A total of 553 units were spike-sorted from control recordings (AP: 44, NTS: 228, DMV: 281) and 459 units from HFD condition (AP: 33, NTS: 192, DMV: 234). Interestingly, the activity of most DVC units varied over the 30 h recording epoch, with a single predominant peak time of firing; these were classified as rhythmic neurons. DVC units that did not exhibit such a peak time in firing were categorised as non-rhythmic neurons. Since short-term consumption of a high-fat diet blunts NTS cFos expression, we anticipated a reduction in the numbers of rhythmic units in the NTS of HFD rats. Unexpectedly, a significantly higher fraction of rhythmic neurons was recorded in the NTS of high-fat diet fed rats (CD: 77.2% vs. HFD: 84.9%, *p*=0.0480, Fisher’s exact test). No diet-related differences in the proportion of rhythmic units were found for the AP (CD: 75% vs. HFD: 84.8%, *p*=0.3974) or DMV (CD: 84.3% vs. HFD: 79.9%, *p*=0.2031).

Manual assessment of the firing rate peak phase revealed that in control diet fed rats, rhythmic AP units elevated their activity at late day/early night with mean peak at pZT12.4 ± 0.01 h (all means calculated as circular means). Interestingly, amongst all rhythmic units, two distinct subpopulations were clearly observed in the NTS: one peaking at late day (30.7% of rhythmic units, pZT10.4 ± 0.005 h) and one at late night at which the majority of rhythmic units peaked (69.3%, pZT22.9 ± 0.005 h; Fig. 5A, left panel). This overt bimodality of the NTS peak distribution was confirmed by statistical assessment (*p*=0.0280, Hartigan’s dip test; Fig. 5B) and reflected in apparent lack of synchrony amongst peaks in each slice (*r*=0.253 ± 0.07; Fig. 5C). Rhythmic units were also recorded from the DMV and these elevated firing at late night (pZT22.3 ± 0.004 h; Fig. 5A,B), with marked synchrony among peaks (*r*=0.697 ± 0.02; Fig. 5C). Thus, in CD animals, rhythmic units in the AP peak at the day-night transition, with NTS rhythmic units showing two peak times of late day and late night, while those of the DMV peak at late night.

**Figure 5.**
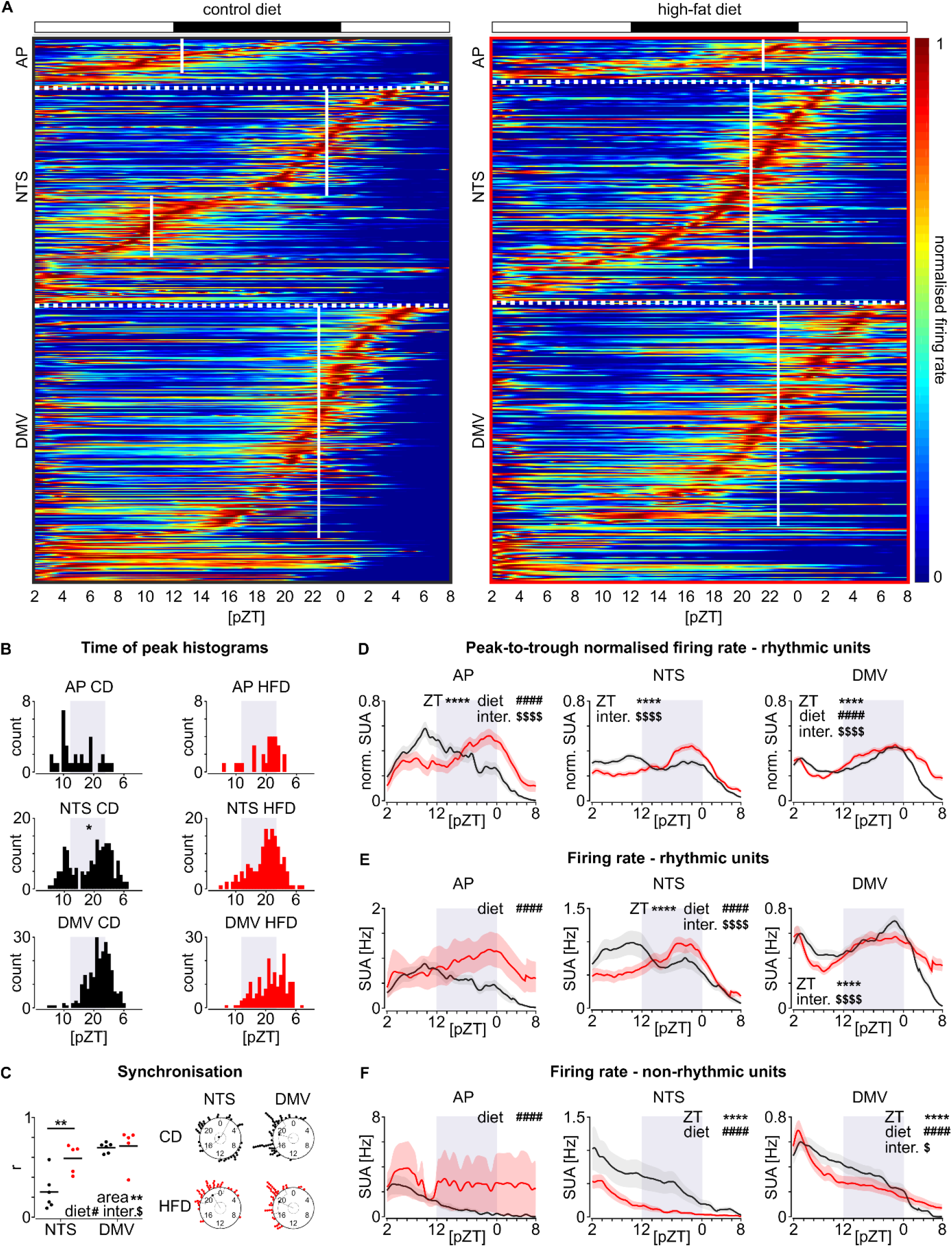
Short-term high-fat diet (HFD) alters the phasing of endogenous circadian firing in the dorsal vagal complex. (**A**) Heatmaps summarising single-unit activity normalised for each unit over the course of 30 h recording of DVC slices (control diet (CD): n=6/563, HFD: n=5/478; slices/units) on the multi-electrode arrays *ex vivo*. Black box surrounds data from CD, whereas red – from HFD. White vertical lines denote circular mean peak phase. Dotted horizontal lines classify units to each DVC area: AP – area postrema, NTS – nucleus of the solitary tract, DMV – dorsal motor nucleus of the vagus. Units are segregated by manually selected predominant peaks from the latest, to earliest. Non-rhythmic units are pulled in the bottom, for each area. (**B**) Histograms of manually selected peaks (bin=1 h). Asterisk codes the significance of Hartigan’s dip test for multi-modality of the distribution (\**p*=0.0280). (**C**) Rayleigh *r* value with example Rayleigh plots showing lack of synchrony in the NTS in CD condition (due to bimodal distribution of peaks; \*\**p*=0.0033, Sidak’s test), but not in HFD (area: \*\**p*=0.0019, diet: #*p*=0.0208, interaction: $*p*=0.0369). (**D**) Mean normalised activity traces (bin=60 s) for the AP, NTS and DMV depicted for rhythmic units only. (**E**) Mean non-normalised firing rate traces showing circadian activity of rhythmic units in three DVC areas. (**F**) Mean non-normalised firing rate traces of non-rhythmic units. All ****, #### and $$$$ code *p*<0.0001 for ZT, diet and interaction, respectively (two-way ANOVA), except for $*p*=0.0409. In all graphs and heatmaps: pZT – projected Zeitgeber time. Black codes CD and red – HFD.

Subsequent manual analysis of recordings carried on brainstem slices collected from high-fat diet fed rats showed abnormal phasing of peak firing in the DVC. Rhythmic units in the AP peaked late at night (pZT21.7 ± 0.01 h), a time clearly phase delayed from recordings made from the AP of CD fed rats (*p*<0.0001, Watson-Williams test; Fig. 5A, STab. 6). This shift in peak distribution caused by HFD significantly changed the circadian waveform of the AP activity (diet: *p*<0.0001, interaction: *p*<0.0001, two-way ANOVA; Fig. 5B,D). Intriguingly, the bimodal distribution of peak firing in the NTS was abolished by HFD (*p*=0.7674, Hartigan’s dip test; Fig. 5A,B), with its rhythmic units forming a single peak around pZT20.9 ± 0.005 h (two-way ANOVA interaction: *p*<0.0001; Fig. 5D). Consequently, synchronisation amongst single units in each slice was significantly higher in HFD compared to CD fed rats (*p*=0.0033, Sidak’s test; Fig. 5C). As the proportion of rhythmic units in the whole NTS population was not lowered by HFD, we hypothesise that a subpopulation of neurons that are ordinarily intrinsically activated to peak at late day is phase-delayed by the high-fat diet such that they achieve peak firing in the late night. Rhythmic units in the DMV of high-fat diet fed rats exhibited maximal firing rates at a similar late night phase as those of CD rats (HFD, pZT22.5 ± 0.005 h; Fig. 5A, right panel). However the shape of the circadian waveform in the DMV was broadened and blunted by diet (diet: *p*<0.0001, interaction: *p*<0.0001; Fig. 5D). The statistical assessment of a possible time shift in the DMV peak activity did not show any significant differences between diets (*p*=0.5813, Watson-Williams test; Fig. 5A, STab. 6).

Further analysis of the absolute, non-normalised firing rates revealed elevated activity in both rhythmic and non-rhythmic units of the AP in HFD rats (diet: *p*<0.001; Fig. 5E,F). In the NTS, high-fat diet reduced the overall level of firing of rhythmic units (diet: *p*<0.0001, interaction: *p*<0.0001 Fig. 5E), in contrast to our observations from the acute protocol. Importantly, the same lowering of the NTS activity was seen in non-rhythmic units (diet: *p*<0.0001; Fig. 5F). In rats fed the high-fat diet, the firing rate of the non-rhythmic DMV subpopulation was also reduced (diet: *p*<0.0001, two-way ANOVAs; Fig. 5F). Further investigations of the possible daily rhythms in the neuronal activities of the DMV and their changes under high-fat diet are presented in our companion study (REF). Collectively, these long-term brainstem recordings indicate pronounced HFD-related alterations in the circadian phasing of neuronal activity among DVC components and highlight the absence of the late day rise of NTS neuronal activity in HFD rats. These findings raise the possibility that the disruption in diurnal feeding pattern arising through consumption of a high-fat diet is underpinned by the suppression of NTS neuronal activity and altered circadian phasing in the brainstem. Statistical summary is presented in Supplementary Table 6.

### 3.5. Daily variation in the responsiveness of NTS neurons to metabolically relevant peptides is altered by HFD

NTS neurons electrophysiologically respond to a variety of orexigenic and anorexigenic signals which modulate their activity level according to the animal’s metabolic status. For example, the peptides orexin and GLP-1 exert potent actions on neuronal activity in the DVC and function as preprandial and postprandial cues, respectively (Peyron *et al*., 1998; Yang & Ferguson, 2003; Yang *et al*., 2003; Richard *et al*., 2015; Fortin *et al*., 2020). Thus, we tested whether responsiveness of NTS neurons to orexin and GLP-1 changes over 24h, and if so, whether this responsiveness is affected by diet. To address this, we used the acute protocol described above to record basal NTS neuronal activity on the MEA platform and then assessed how the amplitude of responses to orexin A (OXA; 200 nM) or GLP-1 (1 µM) (shown as a maximal change in the firing rate) varied at four time points over the day in CD and HFD rats. The majority of NTS neurons responded to OXA (Fig. 6A), whereas GLP-1 evoked responses in less than half of neurons tested (Fig. 6D). The amplitude of response to OXA exhibited time of day variation, but only in the CD condition (*p*=0.0123, two-way ANOVA interaction; Fig. 6B,C), where we observed a clear increase of responsiveness from late day (ZT11) to middle of the night (ZT17; *p*=0.0029). This was not seen in the NTS of HFD rats (*p*=0.9840, Sidak’s tests; Fig. 6B). Time of day variation was also detected in responsiveness to GLP-1 (Fig. 6E,F). Under control diet, NTS neurons elevated their response to this anorexigenic peptide at night (*p*=0.0301, Sidak’s test; Fig. 6E), but in HFD rats, the relation was reversed – the highest amplitude of response to GLP-1 was observed at late day (ZT11) and the lowest at night (ZT17) (*p*=0.0120, Sidak’s test; *p*=0.0123, two-way ANOVA interaction; Fig. 6E). For more statistical details on this dataset see Supplementary Table 5. These findings show that NTS neurons exhibit robust time of day changes in responsiveness to orexigenic and anorectic signals, with this blunted or shifted by high-fat diet.

**Figure 6.**
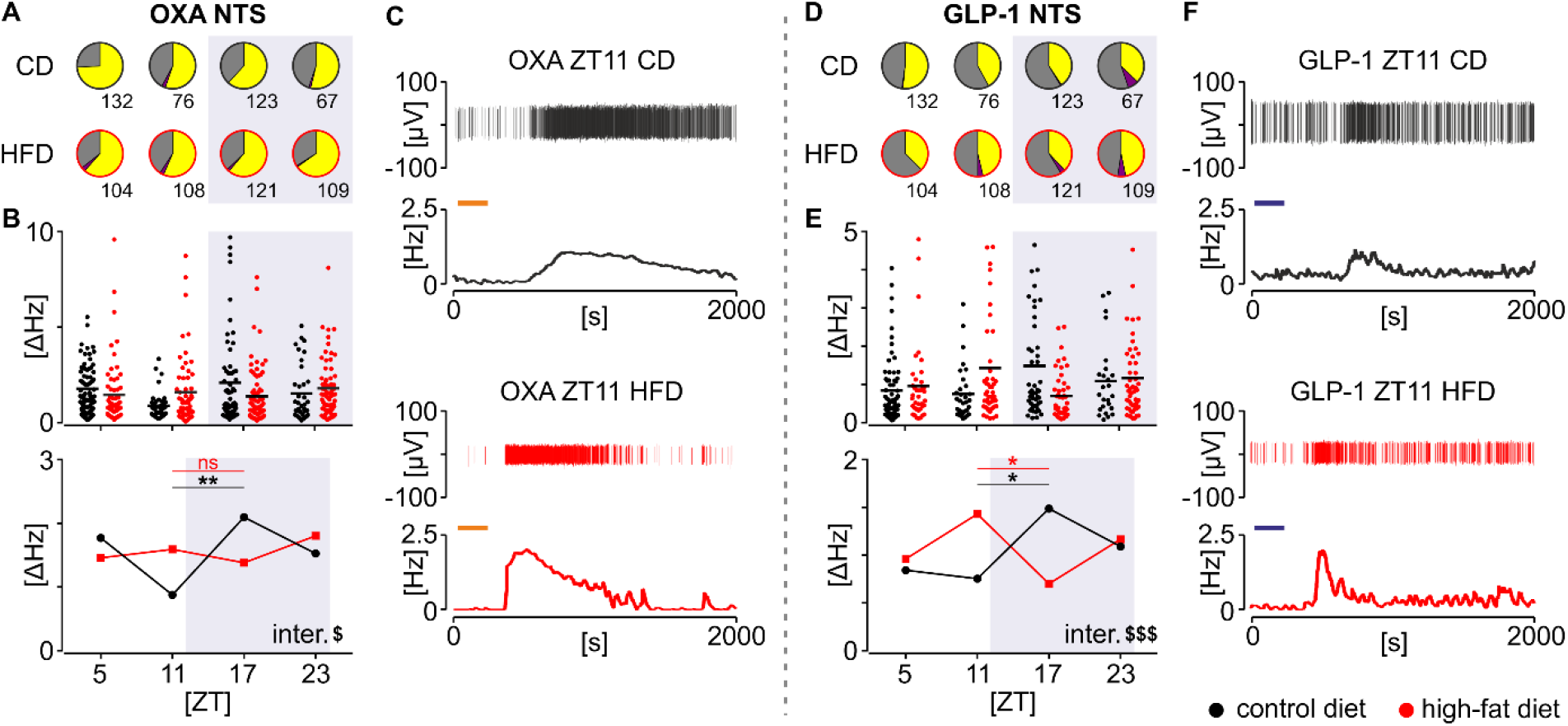
Consumption of a high-fat diet (HFD) alters daily neuronal responsiveness to orexin A (OXA; 200 nM) and glucagon-like peptide-1 (GLP-1; 1 μM) in the nucleus of the solitary tract (NTS). (**A, D**). Pie charts representing the proportion of single units activated (in yellow), inhibited (in purple) and non-responsive (in grey) to OXA and GLP-1 treatment in the NTS tested at four time points across 24h (ZT5, 11, 15 and 23). (**B, E**) Scatterplots (*top*) and summary plots of the mean (*bottom*) showing the amplitude of drug-evoked response. Significant differences between HFD and control diet in the daily pattern of responsiveness to OXA ($*p*=0.0123, two-way ANOVA interaction; CD: \*\**p*=0.0029, HFD: ns *p*=0.9840, Sidak’s test) and GLP-1 ($$$*p*=0.0005, two-way ANOVA interaction; CD: \**p*=0.0301, HFD: \**p*=0.0120, Sidak’s test). (**C, F**) Representative responses of NTS single units to OXA (orange bar) and GLP-1 (blue bar) at ZT11 and 17, presented as spike-sorted single unit recordings and firing rate plots (bin=30s) for both control and HFD. Black codes CD and red – HFD.

## 4. DISCUSSION

Here, we describe time of day and circadian changes in rat NTS neurons and provide substantive evidence of the regulation of DVC neuronal activity by diet. We show that short-term consumption of high-fat diet disturbs the 24h pattern of ingestive activity through elevation of daytime feeding; this is accompanied by misalignment in circadian phasing of DVC neuronal oscillators as well as alteration in the NTS firing rate and responsiveness to metabolic cues. Further, we observe that these neuronal and behavioural changes occur prior to excessive body weight gain. These findings provide new insight into how time of day and diet cues are integrated to alter NTS neuronal activity.

Consumption of a high-fat diet can alter circadian rhythmicity by increasing the period of locomotor activity rhythm. It also attenuates the day-night pattern of feeding behaviour to promote an increase in daytime feeding in nocturnal animals (Kohsaka *et al*., 2007; Furnes *et al*., 2009). We also found that 3-4 weeks of consumption of high-fat diet increased daytime feeding to blunt the day-night rhythm in food intake of adolescent rats. As this occurs prior to the animals gaining excessive weight, the altered time of day/circadian regulation of food intake is not due to non-specific feedback effects of abnormally large body mass, but rather is attributable to consumption of a high-fat diet itself. However, high-fat diet would most likely lead to increased body mass later on, as such trend was observed in HFD rats, but did not reach significant proportions in 3-4 weeks. The DVC is critical for controlling food intake (Grill & Hayes, 2012; Challet, 2019) and rhythmic expression of clock genes in the NTS (Chrobok *et al*., 2020) are attenuated by long-term consumption of a HFD (Kaneko *et al*., 2009). Therefore, here we tested the possibility that HFD disturbs the daily pattern of food intake and that this is accompanied by alterations in the daily and circadian activity profiles of NTS neurons.

First, we evaluated the day-night pattern of cFos expression (a proxy of neuronal activity or increased neuronal input) in the NTS under control and HFD conditions. The number of NTS cells expressing cFos has been tightly correlated with stomach distention (Sabbatini *et al*., 2004) and reported to follow the daily/circadian pattern of feeding (Johnstone *et al*., 2006; Ubaldo-Reyes *et al*., 2017). Therefore, it is difficult to distinguish whether this temporal variation in NTS cFos is attributable to the endogenous circadian activity of this structure or is a consequence of daily changes in neuronal and humoral input arising from the nocturnal pattern of ingestive activity. Our findings indicate that when fed control diet, there is a prominent day to night increase in cFos expression in the NTS which parallels the substantial increase in stomach weight from middle of the day (ZT6) to middle of the night (ZT18). This is preceded by a conspicuous early to late day increase in NTS neuronal activity, as revealed by our long-term MEA recordings. Unexpectedly, cFos levels in the NTS of HFD rats are higher during the day but lower at night, with the peak-nadir amplitude substantially reduced. We posit that this elevated daytime cFos in HFD animals occurs because HFD rats eat more during the day than CD animals. This results in increased stomach distension information conveyed to the NTS; the stomach weight of HFD rats did not differ from ZT6 to ZT18, unlike CD rats. However, we cannot discount the possibility that high-fat diet suppresses NTS neuronal activity prior to the rise in daytime feeding. Alternatively, the reduced night-time levels in the NTS cFos expression of HFD rats may be attributed to the fact that the high-fat diet is calorie dense, thus a smaller chow volume/weight is consumed by animals such that the extent of stomach distension is not as large as that of CD rats.

The temporal variation in cFos *in vivo* suggests that under control diet conditions, NTS neuronal activity increases at night and our acute electrophysiological measurements of the neuronal activity on the MEA *ex vivo* are broadly consistent with this. In the design of our short-term MEA electrophysiological experiments, animals were culled in four time points (ZT3, 9, 15 and 21). It follows then that neuronal activity recorded in the NTS reflects both intrinsic circadian drive as well as feedback from behavioural and physiological processes such as consumption and ingestion preceding the time of cull. Here we show, that under control diet, the NTS activity peaks at late day (ZT11) confirming our recent finding in mice (Chrobok *et al*., 2020) and coinciding with the acrophase in the circadian rhythm of vagal afferent activities (Kentish *et al*., 2016, 2019). This daily variation in CD rats is broadly consistent with rhythm of *Period* clock gene expression in the rodent NTS reported by a number of groups (Herichová *et al*., 2007; Kaneko *et al*., 2009; Chrobok *et al*., 2020; Paul *et al*., 2020). We advance these findings to show that daily/circadian variation in NTS neuronal activities are eliminated in HFD rats; instead NTS neurons from rats fed high-fat diet sustained relatively high levels of activity in the day and notably at late night. This result may suggest a lack of rhythmic peripheral input caused by flattening of diurnal feeding cycle (e.g. constant activation by vagal afferents) or disturbances of intrinsic clock properties of the NTS itself.

To reveal putative endogenous rhythmic properties of circadian oscillators in the DVC, recordings must be carried out in the absence of varying sensory input (e.g. diurnal changes of ambient light or periprandial feedback cues) or input from other circadian oscillators. This experimental approach differs from the acute recordings where animals were culled throughout 24 h or cFos examination *in vivo*, where these activities arise from preceding behavioural state of the animal and its constituting surroundings. Previous long-term recordings (∼24 h) from the mouse DVC *ex vivo* revealed that NTS neurons exhibit pronounced circadian variation in firing rate, increasing to a peak around the late day (ZT10) (Chrobok *et al*., 2020). Since clock genes are rhythmically expressed in the NTS, both *in vivo* and *ex vivo*, our findings raised the possibility that the NTS shares properties with other brain sites, including the SCN, where clock genes drive daily changes in neuronal activity (Guilding *et al*., 2009; Sakhi *et al*., 2014; Takahashi, 2017). Here, in long-term (36 h) recordings of CD rat DVC neuronal activity, we observed a highly organised sequence of peak firing amongst its single units. Notably, two distinct subpopulations of rhythmic units were differentiated in the NTS – one peaking at late day around ZT10, in phase with the upstream AP (as observed previously in mice; Chrobok *et al*., 2020), and the second one elevating its firing at late night hours (around ZT23) at the same phase that some downstream DMV neurons increase firing. Speculatively, this second circadian peak of neuronal activity in the NTS may be explained by a typical pattern in feeding behaviour, which in nocturnal rodents exhibits two night-time peaks – one at the onset of their active phase and the other in advance of the resting day phase (Challet, 2019). Strikingly, the late day elevation in neuronal firing of rhythmic units in the NTS and AP is completely lost in HFD rats, recapitulating our observations from short-term recordings. Since there are no external inputs to the DVC *ex vivo* in these longer term recordings, this suggests that the loss of day-night variation in firing rate is attributable to a high-fat diet altering intrinsic properties of the network of DVC neurons. Thus, a lack of late day peak in HFD NTS may arise from reduced excitatory drive from the upstream AP (whose activity is shifted from late day to late night in this HFD rats) or from local DVC mechanisms disturbed by diet. Interestingly, the discharge of non-rhythmic NTS units was suppressed in high-fat fed animals, suggesting reciprocal inhibition of rhythmic and non-rhythmic NTS cells. Since HFD rats increase daytime feeding to the extent that stomach content does not change from day to night, then this suggests that sustained feedback from stomach distension is responsible for the elevated NTS activity measured from HFD rats in short-term recordings.

Feeding related changes in the DVC are not solely attributable to vagal afferents, but can also arise from afferent input from other brain regions as well as peripheral neurotransmitter/neurohormonal signals (Grill & Hayes, 2012; Challet, 2019). This includes arousal and food-seeking signals from the orexin neurons of the lateral hypothalamus and the anorexigenic cues such as GLP-1 which can originate intrinsically from the NTS as well as extrinsically from gut enteroendocrine L cells (Peyron *et al*., 1998; Crespo *et al*., 2014). Both orexin and GLP-1 systems exhibit circadian rhythmicity (Marston *et al*., 2008; Gil-Lozano *et al*., 2014, 2016; Azeez *et al*., 2018). Intriguingly, loss of orexin neurons or orexin receptors observed in narcolepsy leads to obese phenotype. That is because in healthy individuals orexins not only elicit food intake but promote energy expenditure, which prevents from developing obesity (Yamanaka *et al*., 1999; Hara *et al*., 2001; Funato *et al*., 2009; Kotz *et al*., 2012). Orexin neurons are sensitive to diet and short-term high-fat diet alters the balance of synaptic input to these cells (Linehan *et al*., 2018). Here, we show that the density of OXB-ir fibres in the NTS exhibit time of day variation and under control diet conditions, higher immunoreactivity was found during the day than the night. Since both orexin levels in the cerebrospinal fluid and orexin neuron activation (as indicated by cFos) are lower during the middle of the day than the night (Marston *et al*., 2008; Azeez *et al*., 2018), then the density of OXB-ir most likely reflects the increased accumulation of unreleased peptide during this behaviourally quiescent day phase. Consistent with this interpretation, daytime OXB-ir is reduced in the NTS of HFD rats as these animals engage in active ingestion at this phase of the 24h cycle.

It is notable from the MEA recordings made from DVC slices of control diet fed animals, that NTS neurons show larger responses to OXA during the night, an observation previously reported in mice fed standard laboratory diet (Chrobok *et al*., 2020). Interestingly, this is absent in the NTS of HFD rats where the day-night change in basal NTS neuronal activity is blunted. This suggests that daily variation in NTS neuronal discharge functions to partition enhanced responsiveness to arousal/orexinergic signals to the behaviourally active night. The antiphasic relation between OXB fibre immunoreactivity (presumably indicative of ligand availability) and neuronal responsiveness possibly reflects an active regulatory mechanism whereby NTS neurons increase their sensitivity to orexins at the time of day at which availability of intrinsic orexin peptide is reduced. Unfortunately, the sparseness of intrinsic GLP-1-ir fibres in the DVC (data not shown) precludes similar speculation of ligand availability and neuronal response. As GLP-1 is released from gut enteroendocrine L cells and because the AP lacks a functional blood-brain barrier, then the periphery may be the primary source of this peptide in the NTS. Nearly 50% of rat NTS neurons responded to GLP-1 with the nocturnal increase in responsiveness seen in the DVC slices from control diet fed animals and such increase was absent in slices from HFD rats. Indeed in HFD animals, responsiveness to GLP-1 is maximal during the day, not the night. Interestingly, response magnitude to GLP-1 was negatively correlated with spontaneous firing rates. This may be explained by voltage-dependent mechanisms underlying neuronal response to GLP-1; the lower the membrane potential, the higher the GLP-1-induced excitation (Acuna-Goycolea & van den Pol, 2004). Further patch-clamp investigations are necessary to interrogate synaptic and membrane potential mechanisms underpinning these diet and time of day related responses to orexin and GLP-1.

From our findings, we propose a scenario whereby daily rhythmic changes in the DVC neuronal activity contribute to daily patterns in food intake. We speculate that in standard dietary conditions, the expression of *Period* clock genes drives the AP/NTS neurons to elevate their firing at late day, when the food intake is low, and this serves to maintain an animal in its resting, non-ingesting phase. These endogenous changes are additionally reinforced by higher late day orexin content in axonal terminals in the NTS. Next, at the beginning of the night, the endogenous activity of the NTS begins to decline and the animal initiates ingestion, leading to stomach distension and consequently, a sharp increase in the NTS cFos expression. At the same time, orexin content in the NTS is reduced, plausibly due to increased release during the night. This is paralleled by increased sensitivity of NTS neurons to metabolic cues. Towards the end of the night, animals exhibit anticipatory eating before sleep onset followed by increased neuronal activity of the NTS (the second, late night peak), which occurs as the animal transitions from the behaviourally active to the behaviourally quiescent phase.

This sequence of events dramatically changes under HFD conditions. First, neurons in the AP/NTS lack their endogenous late day increase in spontaneous neuronal activity and this accompanies the elevated daytime consumption of food. We speculate that at this time animals fed HFD do not receive an NTS-derived satiety signal and the homeostatic drive to eat (caused by prolonged rest phase fasting) forces them to consume excessive calories. In response to feedback from daytime feeding, cFos expression in the NTS rises during the light phase and due to the decreased amplitude of daily feeding cycle, it does not exhibit characteristic sharp increase at the beginning of the night. Additionally, since HFD downregulates daytime levels of orexin in the brainstem, activation of the NTS by orexin input during the light phase is blunted. Therefore, we propose that short-term consumption of a high-fat diet alters the daily profile of NTS neuronal activity which over the long-term promotes weight gain and obesity.

## ACKNOWLEDGEMENTS

Authors would like to thank Alan Zmuda, a student at the Department of Neurophysiology and Chronobiology, Jagiellonian University in Krakow, for technical assistance with behavioural procedures. We would also like to thank Patrycjusz Nowik for excellent animal care.

## AUTHORS CONTRIBUTIONS

LC, KPC, JSJL, MHL and HDP conceived the study. LC and MHL supervised the study and provided financial support. LC and JDK designed, performed, analysed and interpreted electrophysiological studies with the help from JSJL. KP wrote custom MatLab and Spike2 scripts for automated spike sorting and further analysis of multi-electrode array data. JDK, LC and JSJL performed spike-sorting. AMS designed behavioural protocols and performed it with LC, KPC and JSJL. AMS and LC analysed and interpreted behavioural study. AMS, KPC, JSJL, JDK and LC performed immunohistochemical studies. LC, JDK and MK performed confocal and epifluorescence imaging. JDK and LC analysed and interpreted histological experiments. LC, JDK and HDP wrote the manuscript and all authors agreed to its final form.

## AVAILABILITY OF DATA AND MATERIALS

The data that support the findings of this study are available from the corresponding author upon reasonable request.

## Abbreviations

ACSF: artificial cerebro-spinal fluid
AP: area postrema
CD: control diet
DMV: dorsal motor nucleus of the vagus
DVC: dorsal vagal complex
FEO: food entrainable oscillator
GLP-1: glucagon-like peptide 1
HFD: high fat diet
MEA: multi-electrode array
NDS: normal donkey serum
NPY: neuropeptide Y
NTS: nucleus of the solitary tract
OXA: orexin A
OXB: orexin B
PBS: phosphate-buffered saline
PFA: paraformaldehyde
pZT: projected ZT
SCN: suprachiasmatic nuclei of the hypothalamus
SD: standard deviation
ZT: Zeitgeber time

## SUPPLEMENTARY TABLES

**Table 1.** Statistical analysis for daily measurements of food intake and body weight throughout four weeks of high fat (HFD, n=10) vs control diet (CD, n=9).

| Body weight (*p*) |  |  |  |  |  |  |  |
| --- | --- | --- | --- | --- | --- | --- | --- |
|  | RM two-way ANOVA |  |  | Sidak's test (CD – HFD) |  |  |  |
| week | ZT | diet | inter. | ZT0 | ZT6 | ZT12 | ZT18 |
| 0 | **<0.0001** | 0.0982 | 0.7197 |  |  |  |  |
| 1 | **<0.0001** | 0.9639 | 0.5830 |  |  |  |  |
| 2 | **<0.0001** | 0.5972 | 0.7501 |  |  |  |  |
| 3 | **<0.0001** | 0.5022 | **0.0110** | 0.9596 | 0.8940 | 0.8995 | 0.9672 |
| 4 | **<0.0001** | 0.1423 | **0.0223** | 0.5014 | 0.3321 | 0.3722 | 0.5054 |
| Food intake (*p*) |  |  |  |  |  |  |  |
|  | RM two-way ANOVA |  |  | Sidak's test (CD – HFD) |  |  |  |
| week | ZT | diet | inter. | ZT0-6 | ZT6-12 | ZT12-18 | ZT18-0 |
| 0 | **<0.0001** | **0.0142** | 0.2663 | 0.9903 | 0.1745 | 0.1991 | 0.2723 |
| 1 | **<0.0001** | **0.0008** | 0.0864 | 0.9817 | 0.5331 | 0.9249 | **0.0007** |
| 2 | **<0.0001** | **0.0002** | 0.7377 | 0.4355 | 0.6049 | **0.0393** | 0.0653 |
| 3 | **<0.0001** | **0.0063** | 0.9369 | 0.5668 | 0.2466 | 0.6380 | 0.8324 |
| 4 | **<0.0001** | **0.0095** | 0.3830 | **0.0369** | 0.9118 | 0.9847 | 0.9964 |
| Body weight at ZT0 (*p*) |  |  |  | Light phase food intake (*p*) |  |  |  |
| RM two-way ANOVA |  | Sidak's test (CD – HFD) |  | RM two-way ANOVA |  | Sidak's test (CD – HFD) |  |
| week | **<0.0001** | week 0 | >0.9999 | week | **<0.0001** | week 0 | 0.9986 |
| diet | 0.5563 | week 1 | >0.9999 | diet | **0.0037** | week 1 | 0.8447 |
| inter. | **0.0003** | week 2 | 0.9920 | inter. | **0.0220** | week 2 | 0.3608 |
|  |  | week 3 | 0.9631 |  |  | week 3 | **0.0153** |
|  |  | week 4 | 0.3978 |  |  | week 4 | **0.0004** |
| Stomach weight (*p*) |  |  |  |  |  |  |  |
| two-way ANOVA |  | Sidak's test (ZT6 – 18) |  |  |  |  |  |
| ZT | **0.0143** | CD | **0.0235** |  |  |  |  |
| diet | 0.3627 | HFD | 0.5541 |  |  |  |  |
| inter. | 0.1830 |  |  |  |  |  |  |

**Table 2.** Statistical analysis for daily expression of cFos in the nucleus of the solitary tract (NTS) in rats fed high fat (HFD, n=24) vs control diet (CD, n=24), with corresponding body weights. N=6 rats/diet/time point.

| Rostral section (*p*) |  |  |  |  |  |
| --- | --- | --- | --- | --- | --- |
| RM two-way ANOVA |  | Sidak's test (HFD) |  | Sidak's test (CD) |  |
| ZT | **<0.0001** | ZT0 vs 6 | 0.4281 | ZT0 vs 6 | 0.7815 |
| diet | 0.0041 | ZT0 vs 12 | **0.0070** | ZT0 vs 12 | **<0.0001** |
| inter. | 0.0083 | ZT0 vs 18 | **0.0022** | ZT0 vs 18 | **<0.0001** |
|  |  | ZT6 vs 12 | 0.4229 | ZT6 vs 12 | **<0.0001** |
|  |  | ZT6 vs 18 | 0.7150 | ZT6 vs 18 | **<0.0001** |
|  |  | ZT12 vs 18 | 0.9989 | ZT12 vs 18 | 0.9999 |
| Intermediate (*p*) |  |  |  |  |  |
| RM two-way ANOVA |  | Sidak's test (HFD) |  | Sidak's test (CD) |  |
| ZT | **<0.0001** | ZT0 vs 6 | **0.0093** | ZT0 vs 6 | 0.9662 |
| diet | 0.0023 | ZT0 vs 12 | **0.0021** | ZT0 vs 12 | **<0.0001** |
| inter. | **<0.0001** | ZT0 vs 18 | **<0.0001** | ZT0 vs 18 | **<0.0001** |
|  |  | ZT6 vs 12 | 0.9966 | ZT6 vs 12 | **<0.0001** |
|  |  | ZT6 vs 18 | 0.5818 | ZT6 vs 18 | **<0.0001** |
|  |  | ZT12 vs 18 | 0.8987 | ZT12 vs 18 | 0.7978 |
| Caudal section (*p*) |  |  |  |  |  |
| RM two-way ANOVA |  | Sidak's test (HFD) |  | Sidak's test (CD) |  |
| ZT | **<0.0001** | ZT0 vs 6 | 0.1045 | ZT0 vs 6 | 0.9477 |
| diet | 0.1169 | ZT0 vs 12 | **0.0001** | ZT0 vs 12 | **<0.0001** |
| inter. | **0.0088** | ZT0 vs 18 | **0.0075** | ZT0 vs 18 | **<0.0001** |
|  |  | ZT6 vs 12 | 0.1269 | ZT6 vs 12 | **<0.0001** |
|  |  | ZT6 vs 18 | 0.9004 | ZT6 vs 18 | **<0.0001** |
|  |  | ZT12 vs 18 | 0.6939 | ZT12 vs 18 | 0.9869 |
| Body weight (*p*) |  |  |  |  |  |
| RM two-way ANOVA |  |  |  |  |  |
| week | **<0.0001** |  |  |  |  |
| diet | 0.7182 |  |  |  |  |
| inter. | 0.1445 |  |  |  |  |

**Table 3.**
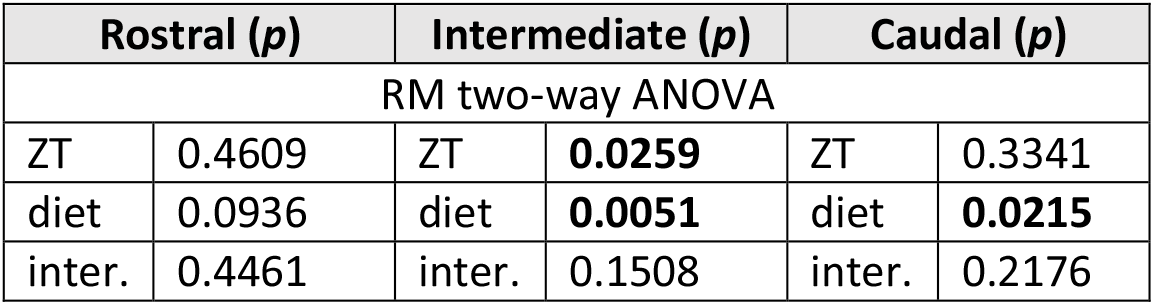
Statistical analysis for daily orexin B (OXB) and neuropeptide Y (NPY)-ir fibres in the nucleus of the solitary tract (NTS) in rats fed high fat (HFD, n=24) vs control diet (CD, n=24), with corresponding body weights. N=6 rats/diet/time point.

**Table 4.** Statistical analysis for short-term recordings of spontaneous neuronal activity in the nucleus of the solitary tract (NTS) with multi-electrode arrays. CD – control diet, HFD – high fat diet.

| NTS (*p*) |  |  |  |  |  |  |  |  |
| --- | --- | --- | --- | --- | --- | --- | --- | --- |
| RM two-way ANOVA |  | Sidak's test (HFD) |  | Sidak's test (CD) |  | n units |  |  |
| ZT | **0.0056** | ZT5 vs 11 | 0.9998 | ZT5 vs 11 | **<0.0001** |  | HFD | CD |
| diet | **0.0001** | ZT5 vs 17 | 0.6871 | ZT5 vs 17 | 0.6915 | ZT5 | 85 | 104 |
| inter. | **<0.0001** | ZT5 vs 23 | **0.0032** | ZT5 vs 23 | 0.9984 | ZT11 | 86 | 111 |
|  |  | ZT11 vs 17 | 0.8849 | ZT11 vs 17 | **0.0091** | ZT17 | 102 | 98 |
|  |  | Z11 vs 23 | **0.0096** | Z11 vs 23 | **0.0012** | ZT23 | 95 | 73 |
|  |  | ZT17 vs 23 | 0.1412 | ZT17 vs 23 | 0.9654 |  |  |  |

**Table 5.** Statistical analysis for short-term recordings of the response amplitude to orexin A (OXA) and glucagon-like peptide 1 (GLP-1) in the dorsal vagal complex with multi-electrode arrays. CD – control diet, DMV – dorsal motor nucleus of the vagus, HFD – high fat diet, NTS – nucleus of the solitary tract.

| OXA NTS (*p*) |  |  |  |  |  |  |  |  |
| --- | --- | --- | --- | --- | --- | --- | --- | --- |
| RM two-way ANOVA |  | Sidak's test (HFD) |  | Sidak's test (CD) |  | n activated / inhibited / total |  |  |
| ZT | 0.1691 | ZT5 vs 11 | 0.9990 | ZT5 vs 11 | **0.0442** |  | HFD | CD |
| diet | 0.9561 | ZT5 vs 17 | >0.9999 | ZT5 vs 17 | 0.8069 | ZT5 | 64/3 /104 | 98/0 /132 |
| inter. | **0.0123** | ZT5 vs 23 | 0.8461 | ZT5 vs 23 | 0.9818 | ZT11 | 61/3 /108 | 42/2 /76 |
|  |  | ZT11 vs 17 | 0.9840 | ZT11 vs 17 | **0.0029** | ZT17 | 74/2 /121 | 76/0 /123 |
|  |  | Z11 vs 23 | 0.9845 | Z11 vs 23 | 0.5169 | ZT23 | 71/1 /109 | 36/1 /67 |
|  |  | ZT17 vs 23 | 0.6434 | ZT17 vs 23 | 0.5342 |  |  |  |
| GLP-1 NTS (*p*) |  |  |  |  |  |  |  |  |
| RM two-way ANOVA |  | Sidak's test (HFD) |  | Sidak's test (CD) |  | n activated / inhibited / total |  |  |
| ZT | 0.5296 | ZT5 vs 11 | 0.2928 | ZT5 vs 11 | 0.9995 |  | HFD | CD |
| diet | 0.8650 | ZT5 vs 17 | 0.8813 | ZT5 vs 17 | **0.0166** | ZT5 | 39/0 /104 | 68/1 /132 |
| inter. | **0.0005** | ZT5 vs 23 | 0.9521 | ZT5 vs 23 | 0.9278 | ZT11 | 50/4 /108 | 32/0 /76 |
|  |  | ZT11 vs 17 | **0.0120** | ZT11 vs 17 | **0.0301** | ZT17 | 46/4 /121 | 50/1 /123 |
|  |  | Z11 vs 23 | 0.8224 | Z11 vs 23 | 0.8507 | ZT23 | 51/6 /109 | 25/5 /67 |
|  |  | ZT17 vs 23 | 0.2474 | ZT17 vs 23 | 0.6502 |  |  |  |

**Table 6.** Statistical analysis for long-term recordings of spontaneous neuronal activity in the dorsal vagal complex with multi-electrode arrays. AP – area postrema, CD – control diet, DMV – dorsal motor nucleus of the vagus, HFD – high fat diet, NTS – nucleus of the solitary tract.

| Time of peak |  |  |  |  |  |
| --- | --- | --- | --- | --- | --- |
|  | CD |  | HFD |  | Watson-Williams test |
|  | ZT | n | ZT | n | *p* |
| AP | 12.4 ± 0.01 | 33 | 21.7 ± 0.01 | 27 | <0.0001 |
| NTS | 10.4 ± 0.004 | 54 | 20.9 ± 0.005 | 163 | <0.0001 |
|  | 22.9 ± 0.005 | 122 |  |  | 0.0002 |
| DMV | 22.3 ± 0.003 | 237 | 22.4 ± 0.005 | 187 | 0.5813 |
| Synchrony - *r* (*p*) |  |  |  |  |  |
| two-way ANOVA |  |  | Sidak's test (CD vs. HFD) |  |  |
| area | 0.0019 |  | NTS | 0.0033 |  |
| diet | 0.0208 |  | DMV | 0.9829 |  |
| inter. | 0.0369 |  |  |  |  |
| Multimodality of peak distribution (Hardigan's dip test) |  |  |  |  |  |
|  | CD |  | HFD |  |  |
|  | *p* | dipStat | *p* | dipStat |  |
| AP | 0.8995 | 0.0454 | 0.9790 | 0.0429 |  |
| NTS | 0.0280 | 0.0416 | 0.7674 | 0.0243 |  |
| DMV | 0.1277 | 0.0303 | 0.2809 | 0.0300 |  |
| Normalised firing rate – rhythmic units (two-way ANOVA *p*) |  |  |  |  |  |
| AP |  | NTS |  | DMV |  |
| ZT | <0.0001 | ZT | <0.0001 | ZT | <0.0001 |
| diet | <0.0001 | diet | 0.6659 | diet | <0.0001 |
| inter. | <0.0001 | inter. | <0.0001 | inter. | <0.0001 |
| Firing rate – rhythmic units (two-way ANOVA *p*) |  |  |  |  |  |
| AP |  | NTS |  | DMV |  |
| ZT | 0.8904 | ZT | <0.0001 | ZT | <0.0001 |
| diet | <0.0001 | diet | <0.0001 | diet | 0.6135 |
| inter. | 0.9635 | inter. | <0.0001 | inter. | <0.0001 |
| Firing rate – non-rhythmic units (two-way ANOVA *p*) |  |  |  |  |  |
| AP |  | NTS |  | DMV |  |
| ZT | 0.9937 | ZT | <0.0001 | ZT | <0.0001 |
| diet | <0.0001 | diet | <0.0001 | diet | <0.0001 |
| inter. | >0.9999 | inter. | >0.9999 | inter. | 0.0409 |

